# MiGut: a scalable *in vitro* platform for simulating the human gut microbiome – development, validation, and simulation of antibiotic induced dysbiosis

**DOI:** 10.1101/2022.09.02.506309

**Authors:** William Davis Birch, Ines Moura, Duncan Ewin, Mark Wilcox, Anthony Buckley, Peter Culmer, Nikil Kapur

## Abstract

*In vitro* models of the human colon have been used extensively in developing understanding of the human gut microbiome and how internal and external factors affect the residing bacterial populations. Such models can be highly predictive of *in vivo* effects of antibiotics, and indeed more so than animal models. The complexity required by current *in vitro* models to closely mimic the physiology of the colon poses practical limits on their scalability. MiGut allows considerable expansion of model runs, increasing the capacity to test reproducibility or parameters under investigation. The MiGut platform has been assessed against a well-studied triple-stage chemostat model in a demanding nine-week study, with exposure to multiple antibiotics inducing a state of dysbiosis in the microbiome. A good correlation is found, both between individual MiGut models and against the triple-stage chemostat. Together with high-throughput molecular techniques for sample analysis, it is now conceivable that tens of *in vitro* models could be run simultaneously, allowing complex microbiome-xenobiotic interactions to be explored in far greater detail. MiGut is a unique platform whereby multiple colonic models can be run simultaneously with minimal added resource and complexity to support our understanding of the cause-effect relationships that govern the gut microbiome. This model system expands the capacity to generate clinically relevant data that can optimize interventions which target the gut microbiome directly or indirectly.

## Introduction

The gut microbiome (GM) is one of the most populous and diverse ecosystems of microorganisms found in the human body^1^, and plays a critical role in host health and disease^2–4^ with disruptions to the intestinal microbiome being correlated with a myriad of health conditions^5, 6^. Historically, studies of the GM have relied heavily on the collection of human fecal samples and the use of animal models to explore the causes and effects of differing microbial compositions of the GM. However, collection of fecal samples can give skewed results^7^, whereas animal and human microbiomes differ in composition and function^8, 9^, limiting their relevance. Additionally, *in vivo* studies require ethical consideration and there is a need to reduce the number of animal species used in research. These limitations can be addressed by *in vitro* models designed to mimic human physiology. They can allow spatial and longitudinal sampling, have fewer ethical constraints, and offer good control over physiological parameters, thus, improving reproducibility.

There exist a range of *in vitro* intestinal models, with varying levels of complexity^10, 11^. The simplest consist of a single stage fermentation vessel which can be batch^12^ or continuously^13^ fed. The relatively low complexity of these models means they can more easily be scaled up: for example, the MiniBioReactor Array (MBRA) can operate up to 48 reactors in parallel to study *C. difficile* physiology and pathogenesis^14^. However, single-stage models lack the spatial bacterial differentiation present in the colon, making them less clinically reflective. By arranging multiple vessels in series, it is possible to simulate different colonic environments – this has been achieved in models such as SHIME^15^ and TIM^16^. A triple-stage chemostat model (supplemental file, section 1) has been extensively used by our team over two decades to investigate the effects of antibiotics on the GM on the large intestine^17, 18^ and the pathogenesis of infections caused, for example, by *Clostridioides difficile* and multi-drug resistant Gram-negative bacteria^19, 20^. This human gut model (HGM) has been validated against human gut content^18^. Although multi-stage models have clear advantages in terms of clinical reflectiveness, the additional complexity means assembly and running of such models is often difficult and resource-consuming, with large costs and space requirements^21^. As a result, experimental replicates are often impractical which limits the complexity of studies that can be performed. There is often a trade-off between process information/control and throughput, illustrated in Figure 1, and there are currently no scalable platforms which adequately simulate the colonic environment.

**Figure 1.**
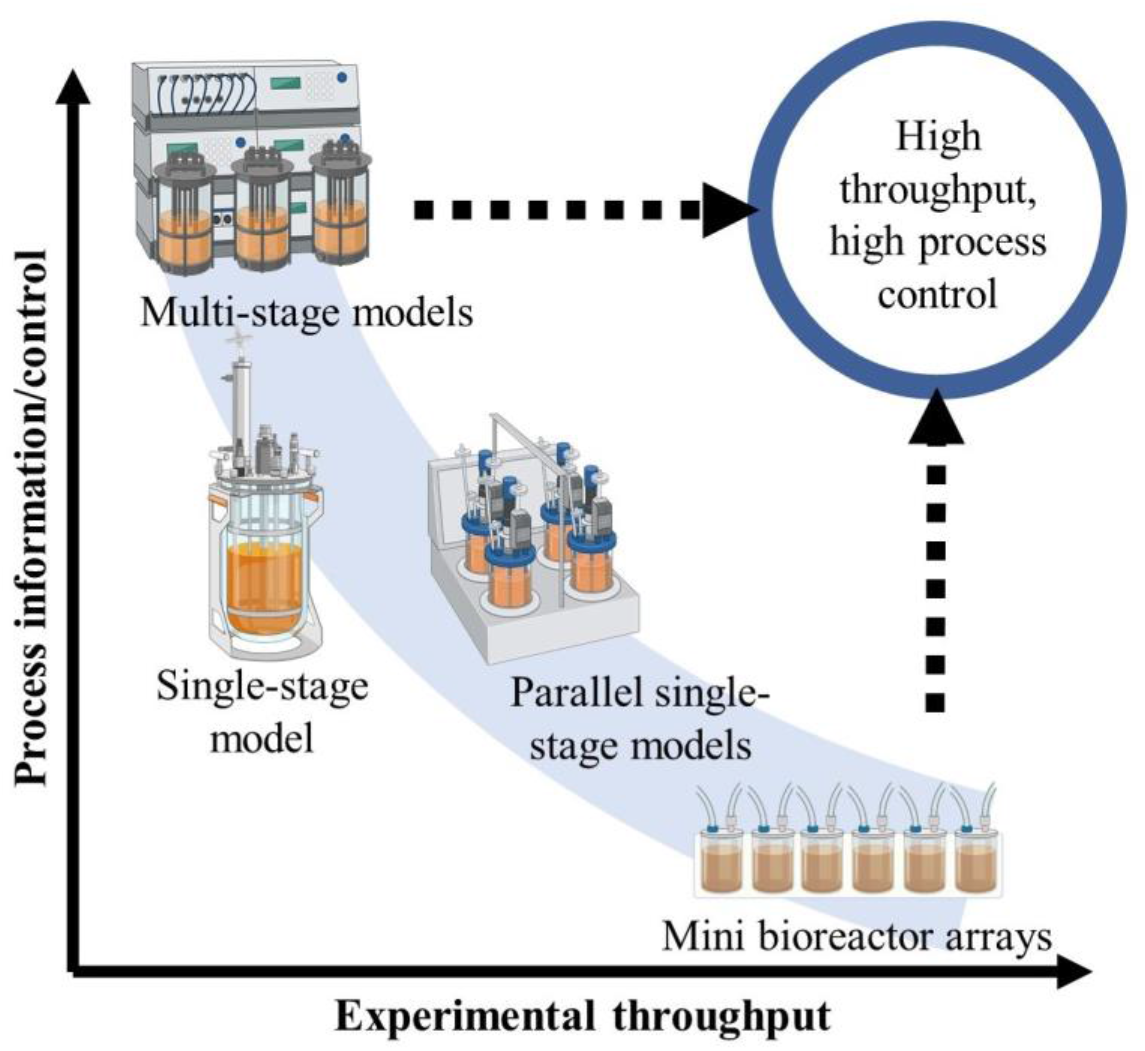
Balance of throughput and depth of process information for a range of bioreactor models. Our work targets the top right quadrant. Created with BioRender.com

An alternative strategy is the use of “gut-on-a-chip” devices, which allow the co-culture of human intestinal cells with the commensal microbiome to study the host-microbe interaction^22, 23^. These microfluidic devices typically contain channels separated by a porous membrane to which either gut epithelial cells or vascular endothelial cells adhere. These are typically perfused with bacteria-free supernatants and lack the control of chemostat models but can be used with more representative fluid taken directly from colonic models, allowing the simulation of bacterial adhesion to the gut wall, for example^24^.

The Mini Gut (MiGut) platform builds upon current *in vitro* technologies to create a scalable colonic simulator with increased experimental throughput compared with other models. Unlike other scalable systems (such as MBRA), MiGut retains clinical relevance by retaining all the functionality of other fully instrumented triple stage models. It takes advantage of recent developments of high-throughput molecular analysis techniques such as real-time PCR to overcome time consuming culture-based methods^25^. To demonstrate the capabilities of MiGut, a nine-week study was performed to show (i) the platform can support a complex human microbiome over an extended period; (ii) the microbial composition within multiple MiGut models behaves consistently when exposed to external stressors, such as multiple antibiotics; (iii) the results of model are directly comparable to those of the clinically reflective HGM.

## Results

### Development of novel platform

An iterative design cycle approach was used to develop MiGut, with outcomes judged against the performance of a well validated, clinically reflective HGM^26, 27^. Acceptance criteria for the model included process conditions of accurate pH (5.5±0.1, 6.25±0.1, 6.75±0.1) and temperature (36.5±0.5°C) across all three vessels, maintenance of an anaerobic environment and physiologically relevant flow rates, in tandem with minimization of day-to-day operational requirements. The engineering process focused on (i) miniaturizing the system to reduce bench space; (ii) user-centric design for easier model setup and operation; (iii) use of Internet of Things technology to improve process control, connectivity, and allow for an information-intensive operation in line with Industry 4.0.

The novel design of MiGut is shown in Figure 2a. Each MiGut platform consists of four triple-stage models, with each vessel representing the proximal (V1), medial (V2), and distal (V3) colon, but containing a much lower volume (45 mL) compared to the HGM (300 mL). The MiGut platform also has a controller, a pumping unit, and a media feed. The models are each machined from a polycarbonate block (Figure 2c), with the internal geometry specifically designed to maintain proper flow direction and promote mixing through the bubbling of nitrogen. Model temperature is maintained with silicone heaters, and the pumping unit controls the pH of each vessel by selectively adding 0.1M HCl or 0.1M NaOH where required. The use of flow selector valves (detailed in Figure 2b) minimizes the number of ancillary pumps needed for pH control with a single pump servicing up to four models. Control is centralized with temperature and pH continually monitored with EasyFerm Plus PHI Arc 120 probes (Hamilton Company, Switzerland). Data is received by the controller and displayed on a touch screen, which also allows user input for monitoring, maintenance, and calibration. All process parameters are streamed every 2 minutes to a database (Microsoft Azure Cosmos DB) for visualization (Microsoft PowerBI) allowing for real-time process monitoring (Figure 3).

**Figure 2.**
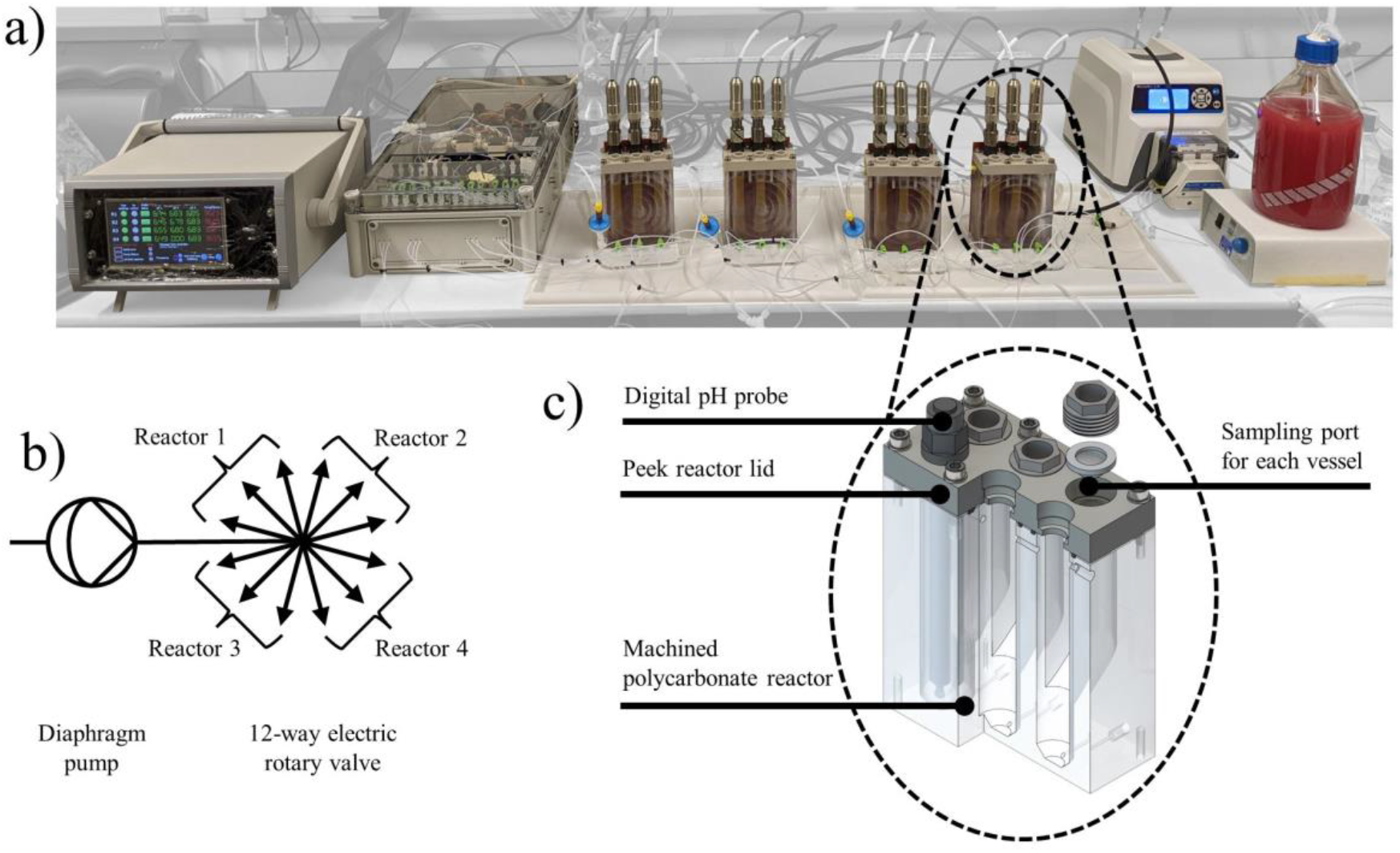
Details of the MiGut model. a) Image of the platform showing the controller, pumping unit, four triple-stage models, and media pump (from left to right) b) Detailed diagram of the acid and alkali pumping schematic for pH control c) Detailed view of the model.

**Figure 3.**
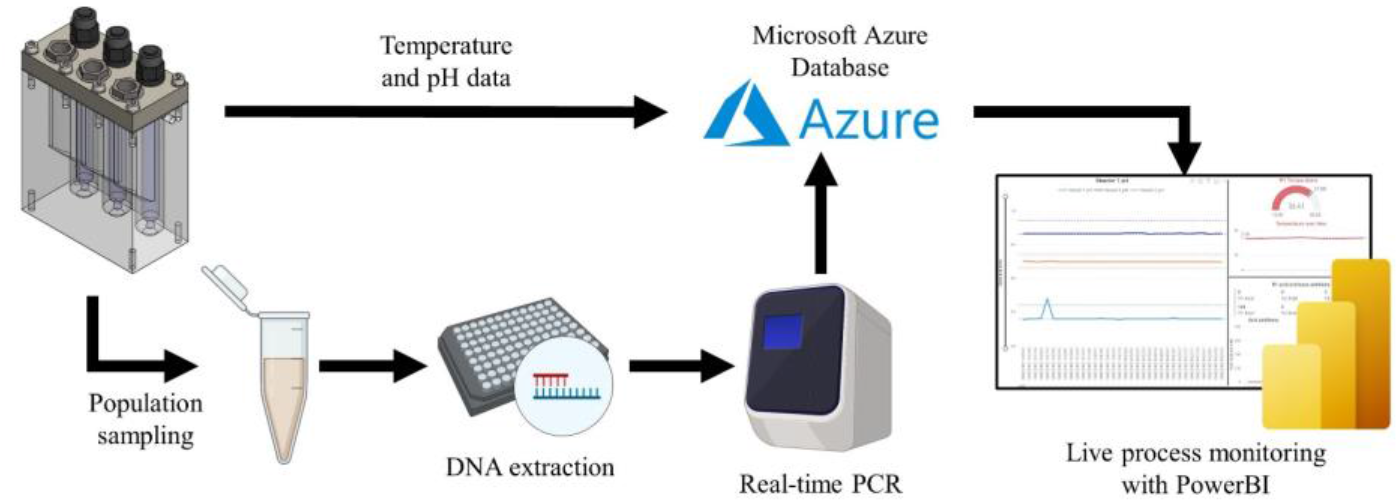
Dataflow for MiGut. Online process data (pH, temperature, acid/base additions) streamed via the cloud for remote process logging and monitoring. Population sampling from PCR giving temporal and spatial sampling. Created with BioRender.com

Due to the reduced vessel volume in MiGut, high-throughput molecular techniques were used instead of culture-based methods to monitor key bacterial populations. It has been shown that real-time PCR correlates well with data from bacterial enumeration when studying *in vitro* models of the GM^25^, with potential for a faster turnaround time than agar sampling. A broad overview of the workflow is shown in Figure 3.

### MiGut forms stable, reproducible communities which correlate with a clinically relevant HGM

The MiGut platform was tested over a nine-week period alongside a HGM, during which the GM was exposed to a series of 3 antibiotics, followed by a four week post antibiotic stage (Table 2). Due to a technical fault in the initial week, one MiGut model was stopped; thus, the data reported are from the remaining three MiGut models.

The initial two-week equilibration stage allows bacterial populations to stabilize in the gut reactor environment, conferring functional properties such as colonization resistance to pathogens. It is often challenging to characterize this community stability: in the HGM, it has been shown that bacterial populations reach an equilibrium and confer functional microbiota properties, such as resistance against pathogen colonization by the end of the two-week period^28^. This was also observed in the MiGut models by using the Bray-Curtis (BC) method, where dissimilarity relative to the fecal inoculum was calculated for each time point during the equilibration stage (Figure 4a). The gradient of each line indicates the rate of change of composition of the microbial communities. All vessels initially had a high rate of change, which plateaued as populations reached an equilibrium (defined by gradient <0.002/day, where the gradient at each point is calculated in three point sliding windows). Based on this analysis, V1 equilibrated on day 6, while V2 and V3 equilibrated on day 8. Upon reaching equilibrium, V1 showed the greatest level of dissimilarity to the fecal inoculum, compared with V2 and V3, whereas V3 was the most similar to the fecal slurry. This is reflective of the microbial populations found within the different sites of the proximal colon (reflected in V1) and the distal colon (reflected in V3). Analysis of similarity (ANOSIM, see Methods) tests showed that the equilibria achieved in V2 and V3 were not significantly different (ANOSIM statistic R (R) < 0.1), while the V1 equilibrium had a statistically significant difference from V2 (R = 0.129) and V3 (R = 0.452). These differences can be visualised by using non-metric multidimensional scaling (NMDS, Figure 4b), where V1 forms a cluster which is largely distinct from those of V2 and V3. Furthermore, the variability in values during equilibrium decreased from V1 through to V3. This is reflected in the standard deviation of the BC dissimilarity (Table 1) and in the decreasing spread of each group in the NMDS plot.

**Table 1.**
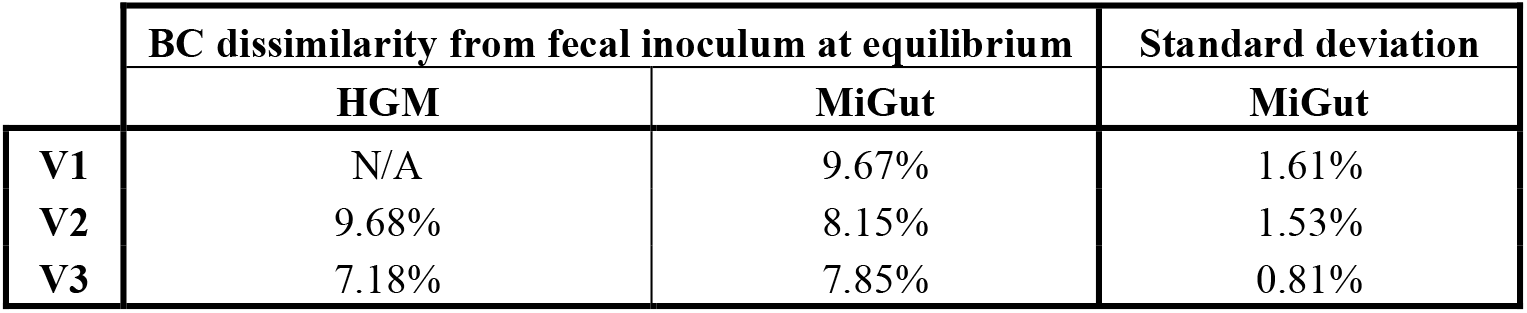
BC dissimilarity data for both the MiGut platform (averaged across all models) and the HGM.

|  | BC dissimilarity from fecal inoculum at equilibrium |  | Standard deviation |
| --- | --- | --- | --- |
|  | HGM | MiGut | MiGut |
| V1 | N/A | 9.67% | 1.61% |
| V2 | 9.68% | 8.15% | 1.53% |
| V3 | 7.18% | 7.85% | 0.81% |

**Figure 4.**
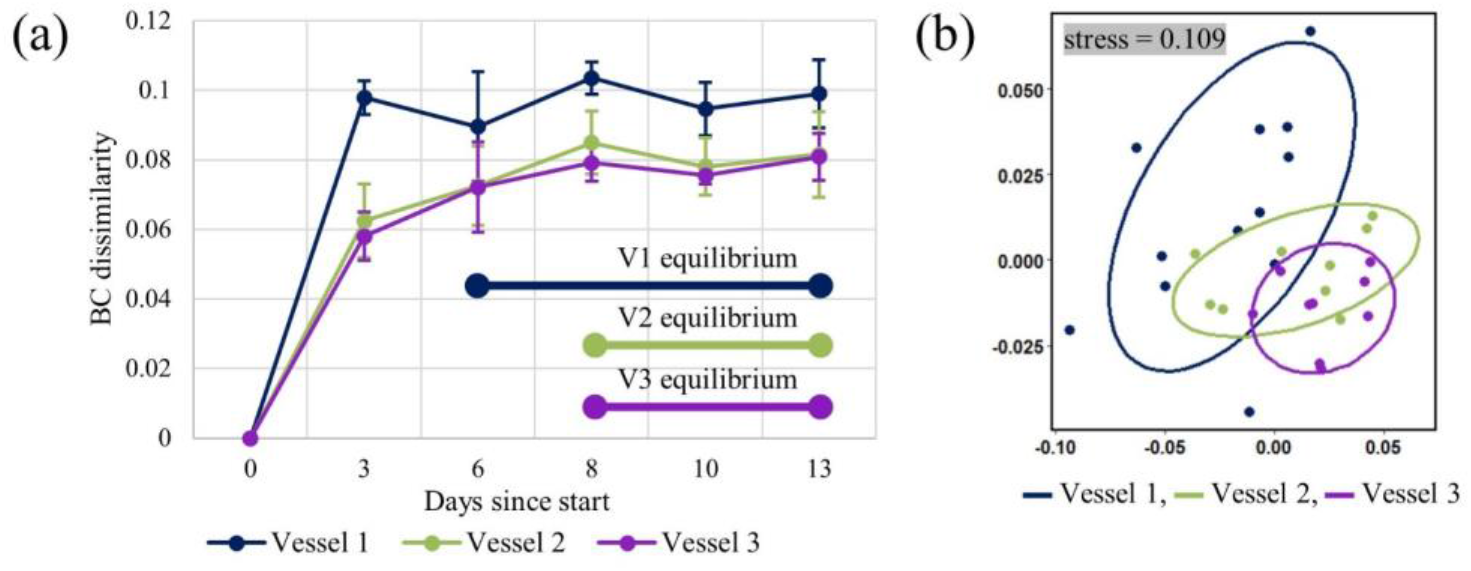
(a) The BC dissimilarity for each MiGut vessel, averaged across all 3 models, during the equilibration period; (b) A NMDS plot of the equilibria of each vessel – each point indicates a sample from the corresponding MiGut vessel (across all models)

The same techniques were used to define equilibrium in the HGM; based on this analysis, the populations in V1 of the HGM did not equilibrate. However, V2 and V3 both equilibrated on day 10, with BC dissimilarities similar to those achieved with MiGut (Table 1). ANOSIM tests were further used to evaluate inter-model similarity (i) between MiGut models, and (ii) between the MiGut platform and the HGM. There was no statistically significant difference between corresponding vessels of the MiGut models (R < 0.1, Figure 5a). When comparing MiGut with the HGM (Figure 5b), V1 and V2 showed high similarity with some differences (0.1 < R < 0.25), while V3 was statistically similar (R < 0.1). Overall, V3 showed the greatest similarity between MiGut and the HGM.

**Figure 5.**
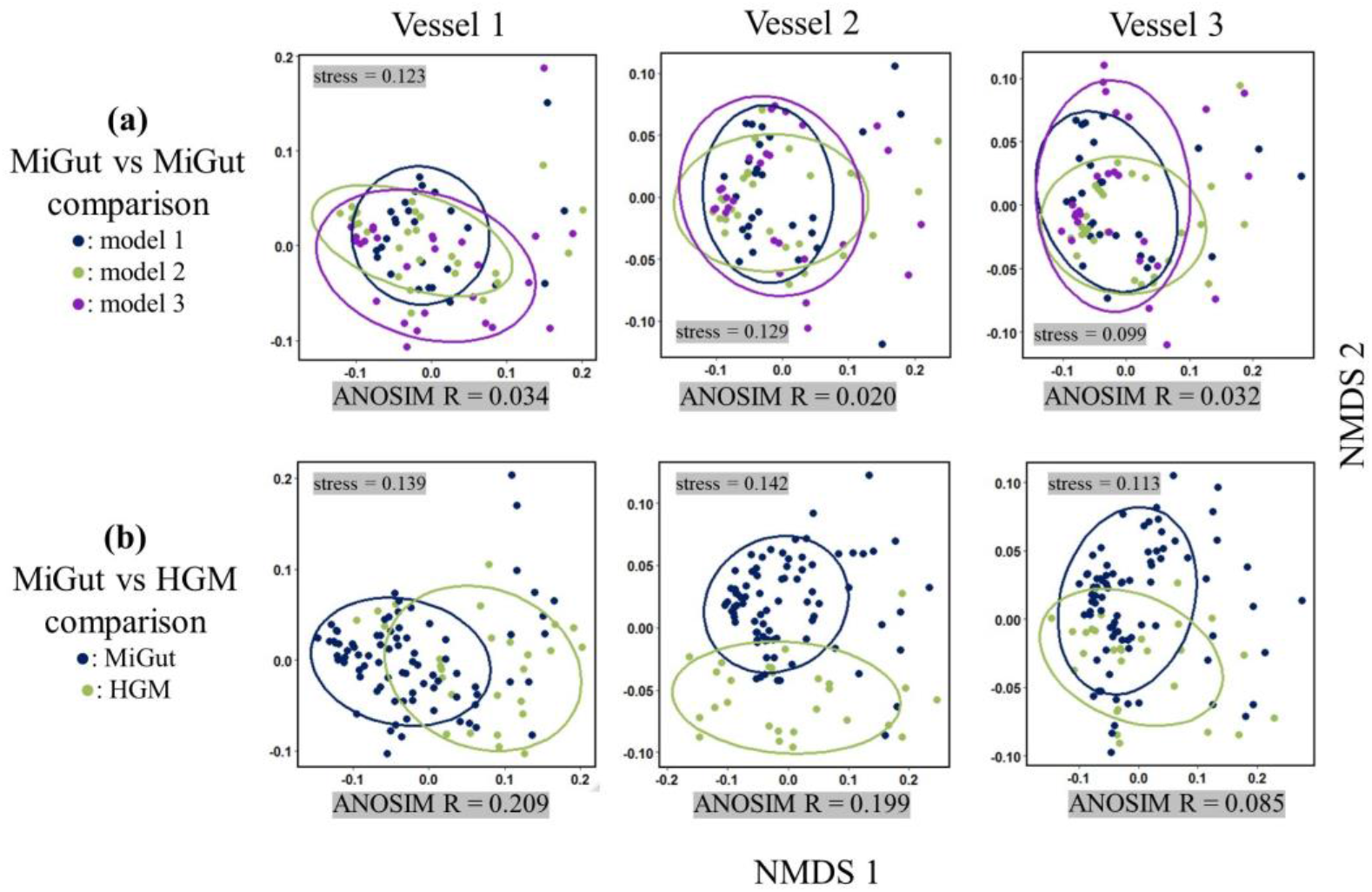
Inter-model comparisons visualised with NMDS plots. (a) shows comparison between MiGut models, (b) shows comparison between MiGut and standard.

Additional BC dissimilarities between MiGut and the HGM were calculated at each stage of the study, shown in Figure 6. Generally, at each stage of the study dissimilarity between systems decreased from V1 being the most dissimilar to V3 being the most similar (note that no comparison is available at the equilibration stage since V1 of the HGM failed to equilibrate). Antibiotic dosing tended to increase dissimilarity and calculated error in all vessels, although the magnitudes here were relatively modest; greatest dissimilarities were 10.8%, 9.5%, and 6.9% for V1, V2, and V3, respectively. The dissimilarity subsequently decreased in the post-antibiotic weeks, with week 4 post-antibiotic having the lowest dissimilarity between MiGut and the HGM for all vessels (4.8%, 3.4%, and 2.4% for V1, V2, and V3, respectively).

**Figure 6.**
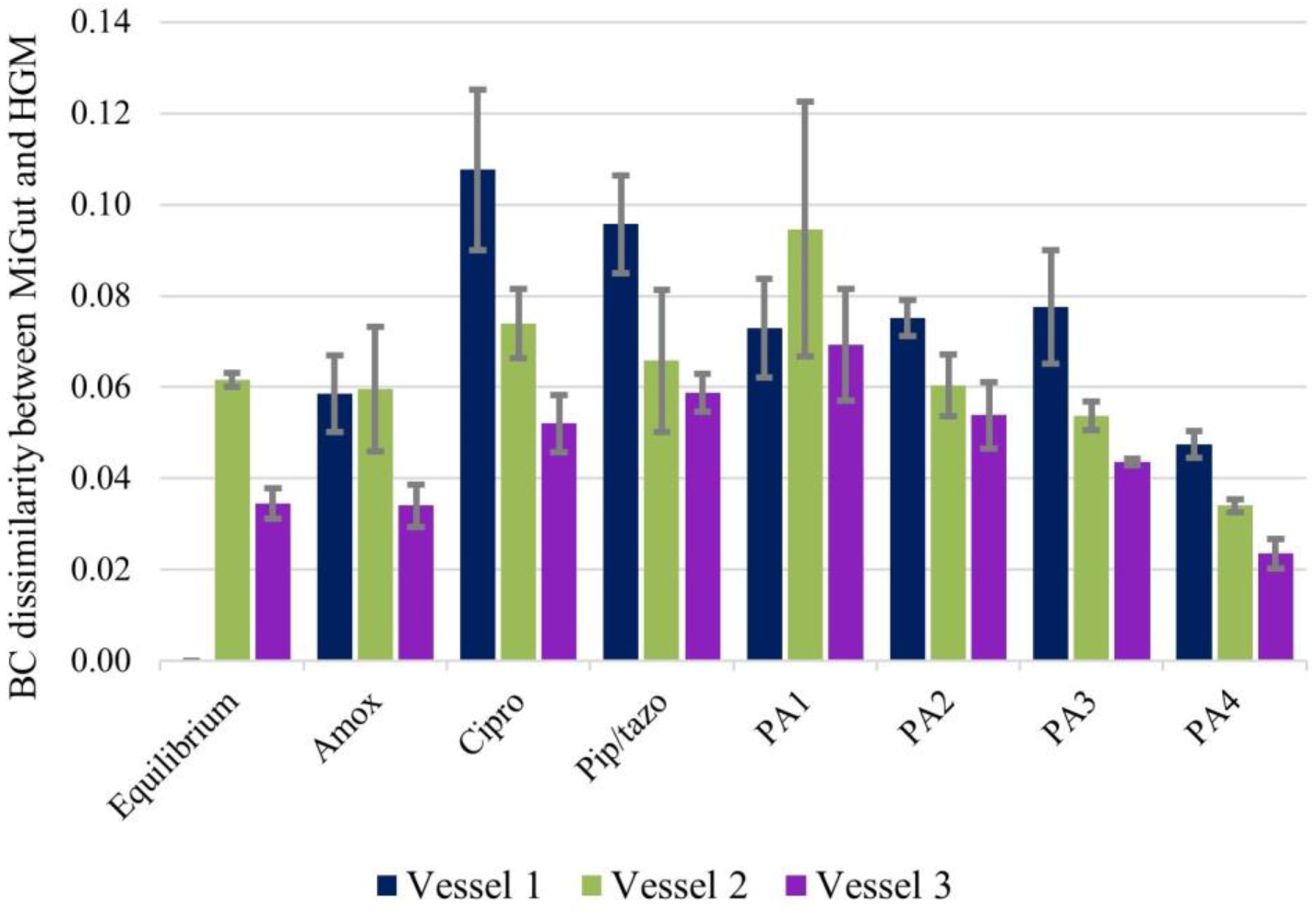
Bray-Curtis dissimilarities between MiGut models (averaged) and the HGM for each stage (equilibrium, amoxicillin, ciprofloxacin, piperacillin/tazobactam, post-antibiotic weeks 1, 2, 3 and 4).

### Antibiotic dosing severely disrupts the microbiome, with MiGut closely matching results from a clinically relevant HGM

Analyses from *in vitro* gut models most commonly focus on V3 which represents the conditions of the distal colon, and so most closely resembles the composition of human fecal samples, allowing for comparison with clinical studies. Accordingly, V3 is considered to be the most relevant when modelling infections^26, 29^ (e.g., *C. difficile* infection). Consequently, we also focused on this vessel when reporting the effects of antibiotic dosing on the residing populations.

BC dissimilarities were calculated relative to the fecal slurry across all stages of the experiment. The results, shown in Figure 7a, indicate that antibiotic dosing disrupted the microbiome from its initial equilibrium, leading to a composition that was overall more dissimilar from the original fecal inoculum. In the subsequent post-antibiotic stages, the microbiome showed signs of recovery as BC dissimilarity once again approached the equilibrium value. However, neither model fully returns to its initial equilibrium, showing that antibiotic dosing has long-term effects. Importantly, the results from MiGut and the HGM correlated extremely well with one another (PMCC > 0.98), with BC dissimilarities being almost identical throughout. The differences in populations’ concentrations between models are explored in detail in Figure 7b. Across the study, the biggest deviation was observed with Bifidobacteria, where the abundance in MiGut was 1.54 log10copies/μL higher than in the HGM. However, this difference subsequently reduced to only 0.31 log10copies/μL. Similar trends were observed in other populations: *C. leptum*, Enterobacteriaceae, and *Lactobacillus spp.*, for example, all show increased dissimilarity during the antibiotic dosing stages. Overall, post-antibiotic week 4 is the stage where MiGut and the HGM are most similar, with a maximum deviation of 0.65 log10copies/μL, also reflected in Figure 7a. Additionally, the product moment correlation coefficient (PMCC) was calculated for each population to quantify how well a change in population in the HGM is replicated in MiGut. Most populations had strong, positive, correlations, indicating that changes in the HGM were reflected in MiGut. The lowest PMCC was that of Enterobacteriaceae (PMCC = 0.65), indicating some differences in the antibiotic response between the HGM and MiGut. Prevotella, which had a high difference in concentration between the models throughout, had the highest PMCC (0.99).

**Figure 7.**
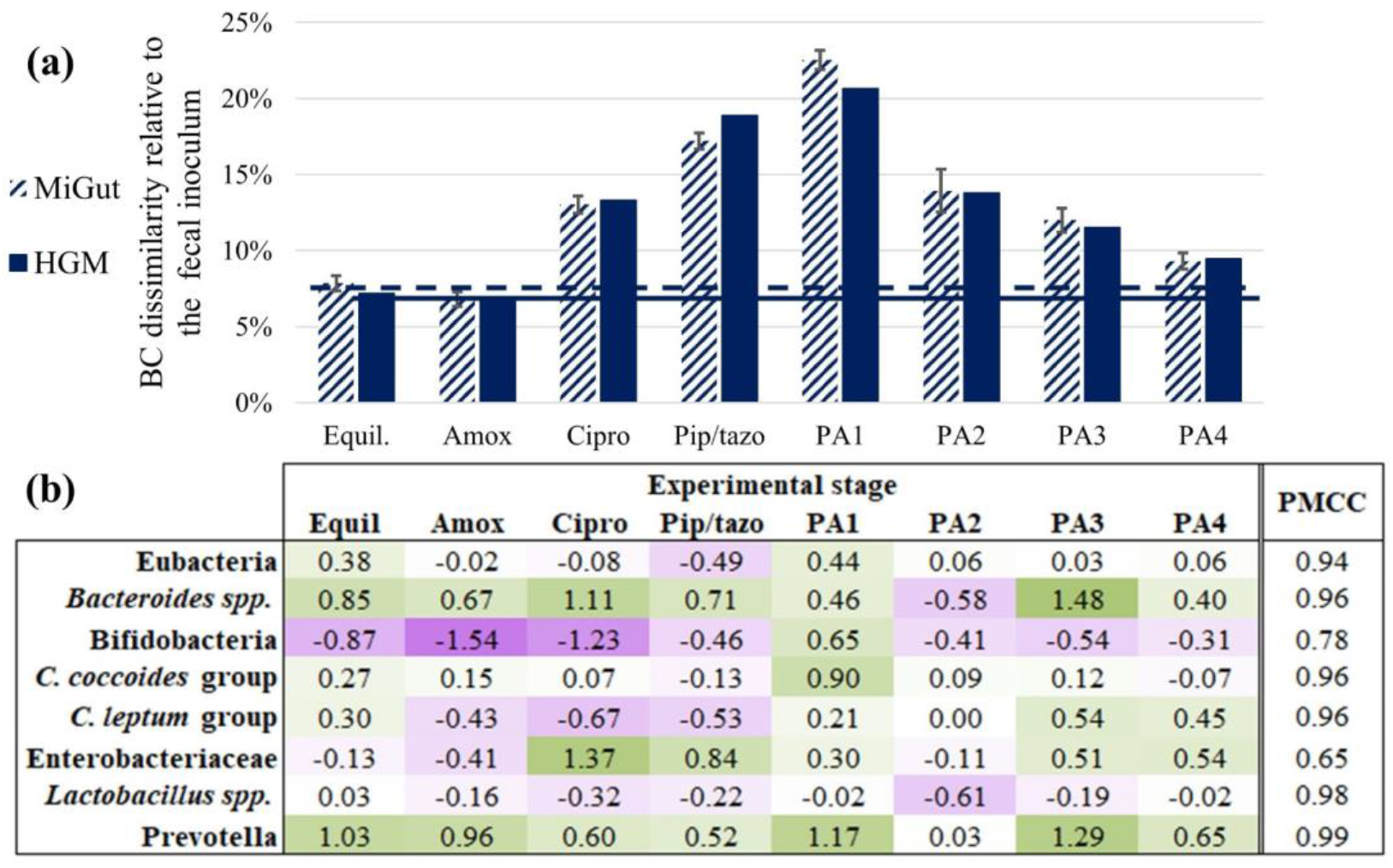
(a) Bray-Curtis dissimilarity relative to the fecal inoculum for V3 of both MiGut and the HGM throughout the study – horizontal lines show the equilibrium value; (b) heatmap showing the difference in (in log10copies/μL) between MiGut and the HGM for each population at each stage – purple cells indicate MiGut > HGM, green cells indicate HGM > MiGut. Additionally, the product moment correlation coefficient was calculated between MiGut and the HGM for each population

Plotting the data on an NMDS plot and grouping by stage (Figure 8) shows that during antibiotic dosing there were pronounced changes in the gut microbiome composition as the groups moved away from the equilibrium region (group 1). This analysis also showed that when the models were challenged with antibiotics, variation increased, reflected by a wider spread of data points. In the post-antibiotic period, when populations were allowed to recover, the variation decreased, and the microbiome composition moves back towards the initial equilibrium. Importantly, the samples in the final post-antibiotic stage (PA4, group 8) formed a closely packed group, which was distinct from that of the initial equilibrium (ANOSIM R = 0.91), indicating that even four weeks after antibiotic withdrawal the bacterial populations did not return to their original state. Within the stage groupings, points corresponding to MiGut and the HGM occupied the same space, showing that there was no significant difference between the two sets of models, further supporting the equivalence of MiGut to the HGM.

**Figure 8.**
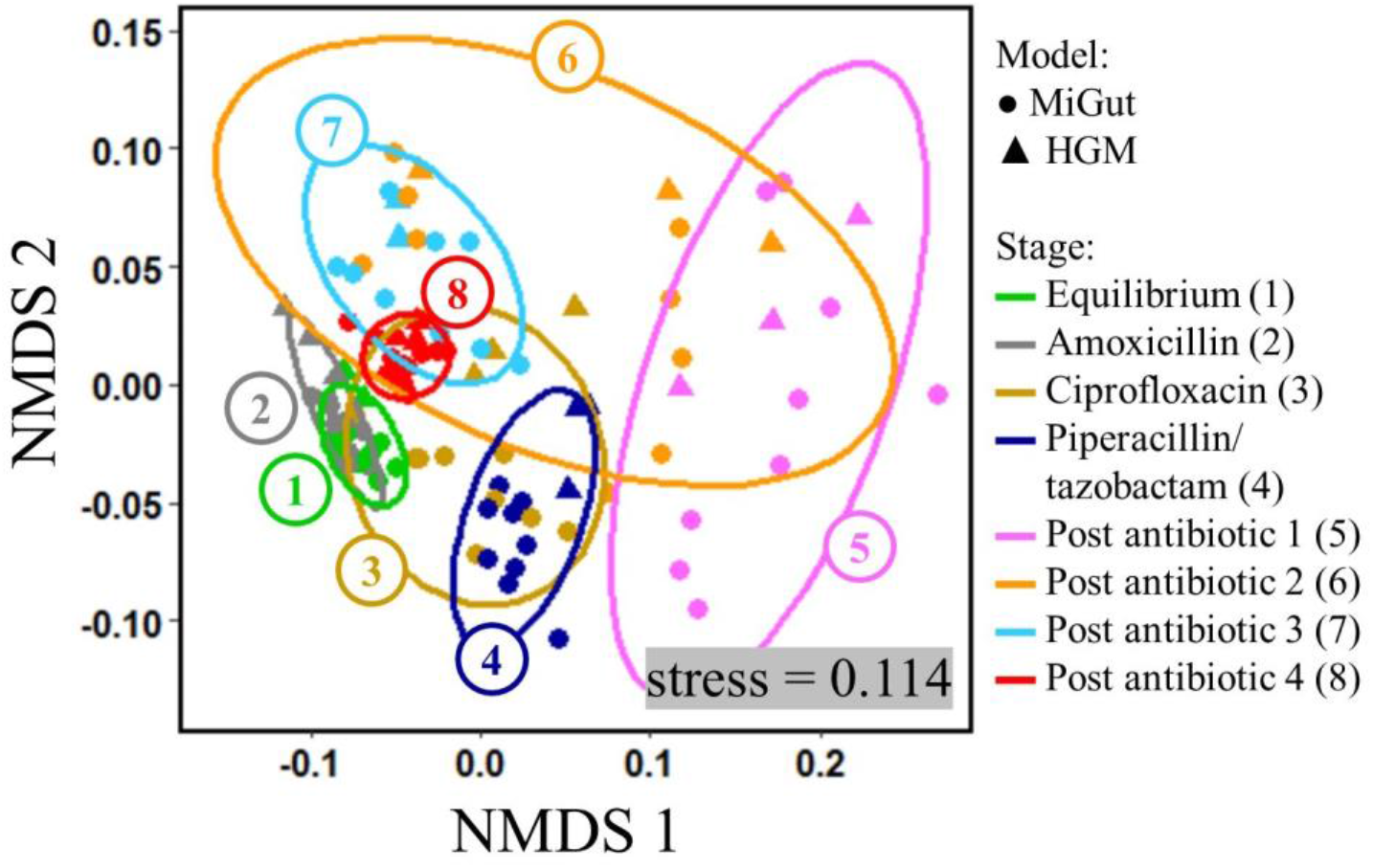
NMDS plot showing the different stages of the experiment from equilibrium through dosing to post antibiotic. Closer grouping of each stage indicates less variability in samples. Groups which overlap with one another have greater similarity than groups which do not overlap at all.

## Discussion

The use of *in vitro* models that aim to simulate the human GM offers a valuable insight into how multiple of internal and external factors can interact to affect the intestinal microbiome. Notably, however, the inherent complexity of these models to closely mimic the colonic environment means that scale-up is often impractical^30^. MiGut has been designed to address these throughput limitations by carefully considering the limiting factors of current technologies and addressing them through an engineering design process. The resulting platform fits four miniature triple-stage bioreactors in the same footprint as one standard gut model^31^. Additionally, the MiGut platform has a reduced setup time and requires minimal user intervention while operational when compared with other models of the human colon. Although this report describes a MiGut platform consisting of four models, it could readily be scaled up to include multiple sets of models running simultaneously in a resource-efficient way.

The MiGut platform was tested in a highly demanding nine-week validation study. During the initial equilibration period, it was important to demonstrate that the bacterial populations were stable across all models. It was found that communities equilibrated within 6 (V1) and 8 (V2 and V3) days of inoculation. The equilibria reached were distinct in V1 compared with V2 and V3, suggesting the physiological conditions of each colonic region were replicated in our triple-stage design. Variability both within and between models during equilibrium was low and decreased from V1 through to V3 (Table 1 and Figure 4a). Furthermore, V1 was more dissimilar to the original fecal inoculum when compared with V2 and V3. This is to be expected since V3 represents the distal colon and is therefore the most comparable with fecal samples. The use of ANOSIM tests (Figure 5a) further demonstrated that the MiGut models showed good repeatability, with no significant differences observed between models.

A single HGM was run alongside the MiGut platform under identical conditions to evaluate the consistency of the new model design compared with the original. Based on the analyses used, V1 of the HGM did not equilibrate in the initial stages and showed a higher degree of dissimilarity between MiGut and the HGM (Figure 5a and Table 1) compared with the other vessels. However, we believe this was due to experimental artefact, where pH fluctuations and volume fluctuations caused some variation of the conditions in the vessel, influencing the microbial populations. This further highlights the benefit of biological replicates. Although there were no replicates of the HGM due to practicality and cost, it has previously been shown that results are consistent between replicates of the HGM^26, 27^.

The decrease in inter- and intra-model variability from V1 through to V3 can be further understood by applying well-known properties of chemical reactors^32, 33^: by considering vessels within a model as well mixed reactors, it can be assumed that an incoming fluid element (nutrient media, antibiotic, acid, alkali) will be instantaneously mixed, displacing a fluid element to the next vessel. This outgoing fluid element will contain a proportion of the incoming element and so on. As a result, the addition of fluid at V1 will cause a large, immediate increase in concentration of that fluid. Subsequently, the outputs from V1 to V2 and from V2 to V3 will have increasingly smoother responses, with lower concentration peaks. Overall, this leads to conditions in V3 being far less variable throughout the study as it is “buffered” by the preceding vessels (supplementary file, section 3).

In this experiment, antibiotic dosing had a profound effect on the GM in V3. Overall, the impact of amoxicillin was less pronounced, but the subsequent antibiotics caused dramatic changes to the bacterial populations, leading to greater dissimilarities compared with the fecal inoculum (Figure 7a). Generally, the concentrations of the measured populations were similar between the HGM and MiGut throughout: *C. coccoides, C. leptum*, and *Lactobacillus spp.* had maximum differences of 0.90, 0.67, and 0.61 log10copies/μL, respectively. Other populations (Bifidobacteria, Enterobacteriaceae, Prevotella) showed more variation. However, by the final week of the study, differences in concentrations for each bacterial population had reduced to 0.65 log10copies/μL or less, indicating that if undisturbed the microbiomes in different models equilibrate to the same levels. The PMCC gives additional insight into how well changes in populations in the HGM are reflected in MiGut: overall, populations correlated extremely well, with 6 out of the 8 reported populations having PMCC > 0.9. Bifidobacteria and Enterobacteriaceae showed some differences (PMCC of 0.78 and 0.65, respectively), mostly during the antibiotic dosing where a greater degree of variability is expected as the populations are subject to external stressors. Interestingly, Prevotella, which had a relatively high concentration difference between MiGut and the HGM throughout, had a PMCC of 0.99: indicating that although there were absolute differences in populations, the relative changes from one stage to another in the HGM were precisely replicated within MiGut.

Overall, these results demonstrate that the MiGut platform can sustain complex bacterial communities with excellent repeatability between models, and provides an environment where communities are able to reliably stabilize. Our study describes the first instance of a scalable platform of fully instrumented triple-stage chemostat models, designed specifically to simulate the human colonic environment *in vitro*. The opportunities that MiGut provides both in terms of reliability of results through biological replicates, and the possibility of studying a far greater number of process variables are truly unique. The human GM is a hugely complex ecosystem, the composition of which can be affected by multiple factors (e.g., age, ethnicity, diet, pathogen exposure, or medication)^2^. MiGut offers the throughput and scalability necessary to investigate the interactions of several factors simultaneously, and to provide assurance data on reproducibility. This will lead to a greater understanding of the cause-effect relationships between xenobiotics and the intestinal microbiome, ultimately informing clinical practice and, we anticipate, leading to more targeted and effective healthcare.

## Materials and methods

### Preparation of inoculum

A pooled fecal slurry was prepared by taking equal samples from five healthy volunteers. Samples were homogenized with phosphate-buffered saline with a 1:5 weight ratio. Each vessel was filled to 50% of its total volume with fecal slurry.

### Setup of the colonic models

Overall, one HGM and one MiGut platform were run in parallel. The HGM was assembled as previously described^25^. Briefly, each model consists of three chemostat vessels of 280 mL, 300 mL and 300 mL, deployed in a weir cascade, top-fed with a complex growth medium at a controlled rate (D=0.015 h^−1^). To reflect the *in vivo* intestinal environment, all vessels were kept at 37°C by a water jacket, continuously stirred, and anaerobically maintained by sparging with N2 to reflect intestinal conditions. The three vessels reflect the pH and nutrient availability observed in the proximal colon (vessel 1, pH 5.5 ± 0.1), medial colon (vessel 2, pH 6.25 ± 0.1), and distal colon (vessel 3, pH 6.75 ± 0.1). Each MiGut model was set up to mimic the HGM. Temperature and pH were identical, and the flow rate of growth medium was scaled to give the same retention time considering the reduction in volume from 300 mL to 45 mL per vessel.

### Experimental Design

All models were inoculated with the same fecal slurry created by pooling human faeces of five donors without history of antibiotic therapy in the previous 3 months. Bacterial populations were allowed to stabilise for 2 weeks (equilibration), and subsequently exposed to antibiotics. Amoxicillin (2 x daily), ciprofloxacin (3 x daily), and piperacillin/tazobactam (2 x daily) were each dosed for five days, with a 2-day recovery/washout period between them, in quantities to give respective concentrations of 24 mg/mL, 278 mg/mL, and 828 mg/mL in V1. Bacterial populations were left undisturbed for a further four weeks after antibiotic dosing. A timeline of the study is shown in Table 2.

**Table 2:**
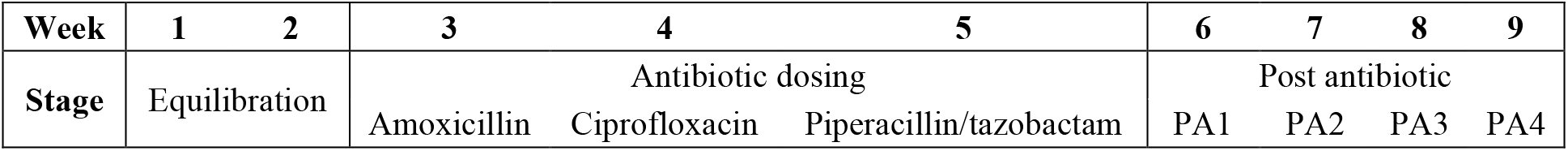
Outline of the nine-week validation study of MiGut.

### Bacterial sampling

Each model was sampled thrice weekly, by collecting 1 mL of fluid from each vessel. Samples were centrifuged (15,000 rpm, 20 min, 4°C), supernatants were removed, and cell pellets were stored at −20^°^C until processing. Following cell disruption using a tissue lyser, DNA extraction was performed using the DNeasy 96 PowerSoil Pro QIAcube HT Kit (Qiagen). DNA extracts were normalized to 5 ng/μL, and 8 key bacterial populations were monitored by real-time as previously described^25^.

### Data analysis

All real-time PCR samples were analyzed against a plasmid curve to calculate bacterial concentrations. Data was converted to a logarithmic scale to allow for normal distribution^25^. Non-metric multidimensional scaling^34^ was used to represent the data in 2D plots. The fit of the NMDS ordination is evaluated with “stress”^35^. Stress < 0.2 is considered an acceptable representation. Analysis of similarities (ANOSIM) uses a dissimilarity matrix to evaluate differences between populations^34^. The generated ANOSIM statistic R gives a measure of significance: R < 0.1 = similar; 0.1 < R < 0.25 = similar with some differences; 0.25 < R < 0.5 = different with some overlap; R > 0.75 different^36^. Bray-Curtis dissimilarity is an ecological measure of dissimilarity between two groups ranging from 0 (identical) to 1 (completely dissimilar)^37^. NMDS plots, ANOSIM tests, and Bray-Curtis dissimilarities were all calculated with RStudio (RStudio: Integrated Development Environment for R, version 2021.9.2.382, www.rstudio.com). Heatmaps (Figure 7b) and PMCCs were calculated with Microsoft Excel (Microsoft Corporation, version 2021).

### Ethics statement

The collection and use of human feces in our gut model has been approved by the School of Medicine Research Ethics Committee, University of Leeds (MREC 15-070 – Investigation of the Interplay between Commensal Intestinal Organisms and Pathogenic Bacteria). Participants were provided with a ‘Participant Information Sheet’ (PIS) detailing a lay summary of the in vitro gut model and the scientific work they were contributing to by providing a fecal donation. Within this PIS, it is explained that by providing the sample, the participant is giving informed consent for that sample to be used in the gut model.

## Acknowledgments

The authors thank Georgina Davis and Vikki Wilkinson for the administrative and material support, and Graham Brown for manufacturing services.

## Authors’ contributions

WDB (William Davis Birch), NK (Nikil Kapur), and PC (Peter Culmer) designed and developed the MiGut platform. AB (Anthony Buckley), IM (Ines Moura), and MW (Mark Wilcox) designed and overviewed the experimental study. WDB, DE (Duncan Ewin), IM, and AB conducted the experiment. IM performed DNA extraction and bacterial quantification. Data was analyzed by WDB. All authors contributed to and approved the manuscript.

## Availability of data and materials

Additional data is available from the University of Leeds at https://doi.org/10.5518/1166

