## Supplemental file for "MiGut: a scalable *in vitro* platform for simulating the human gut microbiome – development, validation, and simulation of antibiotic induced dysbiosis"

Provide short biographical notes on all contributors here if the journal requires them.

### 1. A triple-stage model of the human gut microbiome[1]

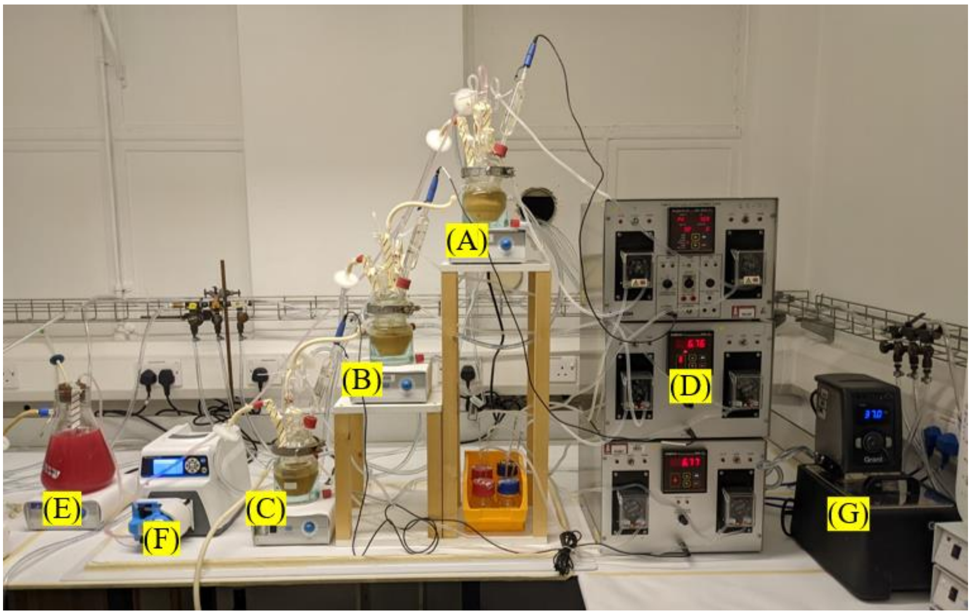


Figure 1: Typical setup of a triple-stage human gut model. (A) Vessel 1, (B) Vessel 2, (C) Vessel 3, (D) pH controllers (one for each vessel), (E) Nutrient-rich media, (F) Peristaltic pump for media, (G) Heated water pump

- Vessel 1 (proximal colon): high nutrient availability, pH 5.4 – 5.6, 280ml
- Vessel 2 (medial colon): low nutrient availability, pH 6.15 – 6.35, 300ml
- Vessel 3 (distal colon): low nutrient availability, pH 6.65 – 6.85, 300ml

Vessel 1 is top fed with a complex growth medium[2] at a constant rate of 0.3 ml/min (simulates a 48h retention time). All vessels are continually sparged with nitrogen.

### 2. Results of the standard triple-stage human gut model

Full Bray-Curtis dissimilarity results are available from the University of Leeds at

https://doi.org/10.5518/1166

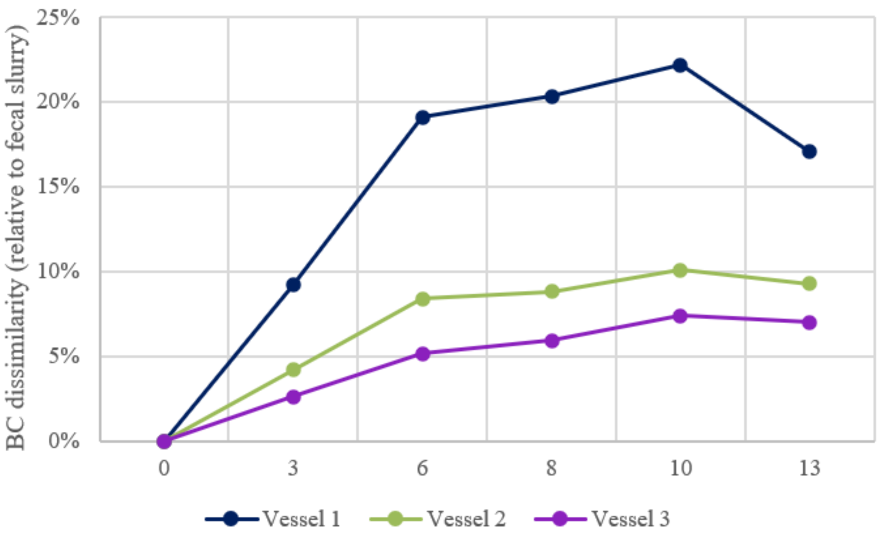


Figure 2: Bray-Curtis dissimilarities from the fecal inoculum for the human gut model during equilibration

#### 3. Variation of input concentrations for each vessel

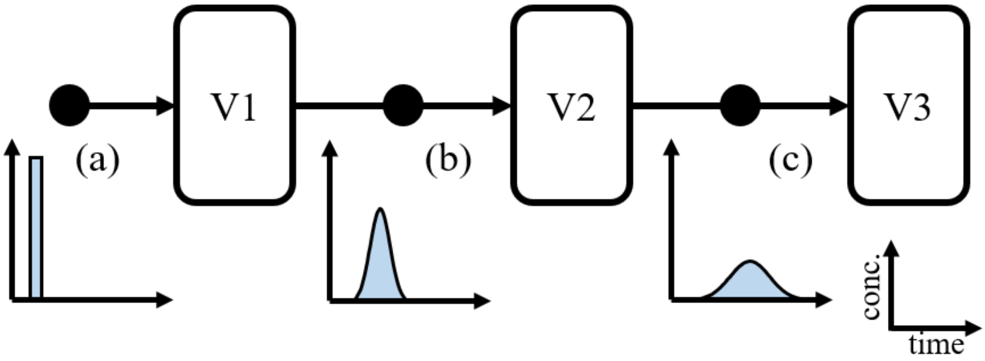


Figure 3: Concentration-time diagrams for each of the inputs to V1 (a), V2 (b), and V3 (c). The shaded area under each graph, representing fluid volume, is identical.
